# An energy-dependent integrate-and-fire model explains coordinated synaptic and intrinsic adaptation under metabolic stress

**DOI:** 10.64898/2026.10.28.757140

**Authors:** Philip Sommer, Peter Jedlicka, Alexander D. Bird, Jochen Triesch

**Affiliations:** Frankfurt Institute for Advanced Studies, Frankfurt am Main, Germany; Department of Physiology, Anatomy and Genetics, University of Oxford, Oxford, United Kingdom; Computer-Based Modelling in the field of 3R Animal Protection, ICAR3R, Faculty of Medicine, Justus Liebig University, Giessen, Germany; Translational Neuroscience Network Giessen, Germany; Institute of Clinical Neuroanatomy, Neuroscience Center, Goethe University, Frankfurt; Ernst Strüngmann Institute (ESI) for Neuroscience in cooperation with the Max Planck Society, Frankfurt am Main, Germany

**Author notes:** Joint senior authors (Peter Jedlicka, Alexander D. Bird, Jochen Triesch).

**Keywords:** neuronal diversity, neuronal energy budget, energy-efficiency, control theory, degeneracy, food restriction, synaptic scaling

## Abstract

Neuronal computation is constrained by the energetic cost of maintaining synaptic transmission, ionic gradients and action-potential generation. Prior work has shown that food restriction in mice shifts visual cortical neurons toward a different operating regime characterized by reduced excitatory synaptic drive, increased membrane resistance and a more depolarized resting membrane potential. This preserves neuronal firing rates, but comes at the cost of reduced coding precision. Deriving an improved estimate of neuronal energy consumption, we have recently shown that these adaptations reflect an optimal trade-off between neuronal information transmission and energy consumption. However, it remains unclear through which control strategies individual neurons adapt their biophysical parameters to maintain a favorable energy-information trade-off. Here we consider several candidate models implementing different control strategies. A „synapses-first” model, in which metabolic stress drives a downscaling of synaptic efficacies to save energy, while an intrinsic plasticity mechanism increases excitability to preserve the average firing rate, successfully reproduces key experimental observations. Alternative models in which energy shortages drive changes in intrinsic excitability or firing-rate variability can also reproduce the endpoint of neuronal adaptation, but predict different time courses and exhibit less energy-efficient and less stable dynamics. New experiments are needed to distinguish among these competing models.

**In brief:**

- A dynamic energy-dependent integrate-and-fire model reproduces food-restriction-like parameter shifts when metabolic feedback first downscales synaptic efficacy and activity feedback subsequently increases intrinsic excitability.
- Reversing this control architecture, so that energy acts first on intrinsic excitability, produces suboptimal adaptation dynamics with transient hyperactivity, making it a less parsimonious explanation.
- Extending this control framework to large-scale recurrent network models could show how prolonged energy disruption destabilizes excitability and contributes to neurological disorders.

**Statement of significance:** Neurons in the brain must function with a limited supply of energy. Recent work has shown that during energy scarcity neurons react by lowering their energy consumption at the expense of reduced information transmission. But through which mechanisms do they regulate their energy consumption and manage to remain as informative as possible? Here, we present a computational model that explains how neurons make coordinated adaptations of their synapses and cell membranes to regulate the trade-off between energy consumption and information transmission. Our work suggests that neurons are efficient energy savers that carefully regulate their energy consumption in response to current energy supply to transmit the maximum amount of information.

## Introduction

Neuronal computation is constrained by energy. Although the brain represents only a small fraction of body mass, it consumes a disproportionate share of the body’s energy budget (Wang *et al*., 2010). This constraint has shaped brain size, wiring, synaptic transmission, neural coding, and information processing (Levy and Baxter, 1996; Schreiber et al., 2002; Chen *et al*., 2006; Niven and Laughlin, 2008; Navarrete *et al*., 2011; Isler and Schaik, 2009; Hyder *et al*., 2013; Harris *et al*., 2015; Yu and Yu, 2017; Harris *et al*., 2019; Padamsey and Rochefort, 2023). Energy availability is therefore not merely a background limitation, but a regulatory constraint that shapes how neurons set their biophysical parameters.

Recent food-restriction experiments in mouse visual cortex provide direct evidence for such energy-dependent adaptation. Under reduced metabolic supply, layer 2/3 pyramidal neurons reduce excitatory synaptic efficacy while increasing intrinsic excitability through higher membrane resistance and a depolarized resting potential (Padamsey *et al*., 2022). These coordinated changes reduce costly synaptic drive while preserving firing rates. However, they also degrade sensory coding, indicating that neurons may trade coding precision for energy savings under metabolic stress (Padamsey *et al*., 2022). While these experiments reveal the endpoint of adaptation, the dynamic rules that coordinate synaptic and intrinsic changes over time remain an open question.

Homeostatic plasticity provides a natural control-theoretic framework for this problem. From this perspective, neuronal activity acts as a feedback signal that is compared with a target regime, and deviations from this target drive compensatory changes in synaptic or intrinsic parameters (Drion *et al*., 2015; Yang and Prescott, 2023). Such negative feedback loops can stabilize neuronal dynamics despite perturbations, parameter fluctuations, and changing input statistics (Niemeyer *et al*., 2021). However, most models of homeostatic activity regulation treat activity as the only control variable and do not include regulation of neuronal energy consumption. This raises a central question: how can a neuron reduce energy consumption while maintaining functional output? Existing energy-dependent integrate-and-fire models have begun to link excitability to energetic variables, but typically regulate only a restricted subset of parameters, such as resting (Fardet and Levina, 2020; Dadras and Park, 2026) or reset potential (Jaras *et al*., 2021). They therefore do not capture the experimentally observed combination of synaptic downscaling, membrane-resistance increase, and resting-potential depolarization induced by food restriction (Padamsey *et al*.,

Here, we introduce an energy-dependent integrate-and-fire framework that links metabolic state to experimentally observed synaptic and intrinsic adaptations. The model combines an updated neuronal energy budget with leaky integrate-and-fire (Brunel and Van Rossum, 2007) and adaptive exponential integrate-and-fire (Brette and Gerstner, 2005) dynamics, allowing the neuron to regulate synaptic strength, resting potential, and membrane resistance. Its simplicity makes it computationally efficient enough for systematic parameter exploration, while its experimentally grounded control rules allow us to compare competing computational mechanisms of energy-dependent adaptation.

We consider three control architectures. In a *synapse-first* model, metabolic feedback reduces ATP-expensive synaptic drive, and activity-dependent intrinsic plasticity compensates to preserve firing rate. In an *excitability-first* model, energy acts first on intrinsic excitability, and synaptic scaling restores activity secondarily. In a *variance-control* model, membrane resistance is regulated by spiking variability rather than mean firing rate. We show that the *synapse-first* architecture most robustly reproduces food-restriction-like parameter shifts, whereas alternative architectures produce less stable dynamics. Our results provide a first computational account of how neurons simultaneously regulate their energy consumption and firing activity to maintain energy-efficient information processing and communication.

## Results

Experimental work has shown that food restriction shifts visual cortical neurons toward parameter configurations that reduce ATP consumption at the expense of coding precision (Padamsey *et al*., 2022). Although this establishes the optimal target of adaptation in parameter space, it leaves open how coordinated synaptic and intrinsic changes emerge dynamically. We therefore formulate a minimal control architecture applied to a phenomenological energy-dependent integrate-and-fire model in which synaptic efficacy and intrinsic excitability are regulated by separate but interacting feedback signals.

The aim of this model is not to reproduce the molecular complexity of metabolic signaling. Instead, it captures the control logic inferred from the experimental data (Padamsey *et al*., 2022, 2024): reduced energy availability downscales ATP-expensive excitatory synaptic drive, whereas activity-dependent homeostatic plasticity adjusts intrinsic excitability to stabilize neuronal firing within a functional range (Fig. 1). This formulation provides a compact data-driven framework for studying how energetic constraints can shape neuronal parameters over long time scales.

**Figure 1:**
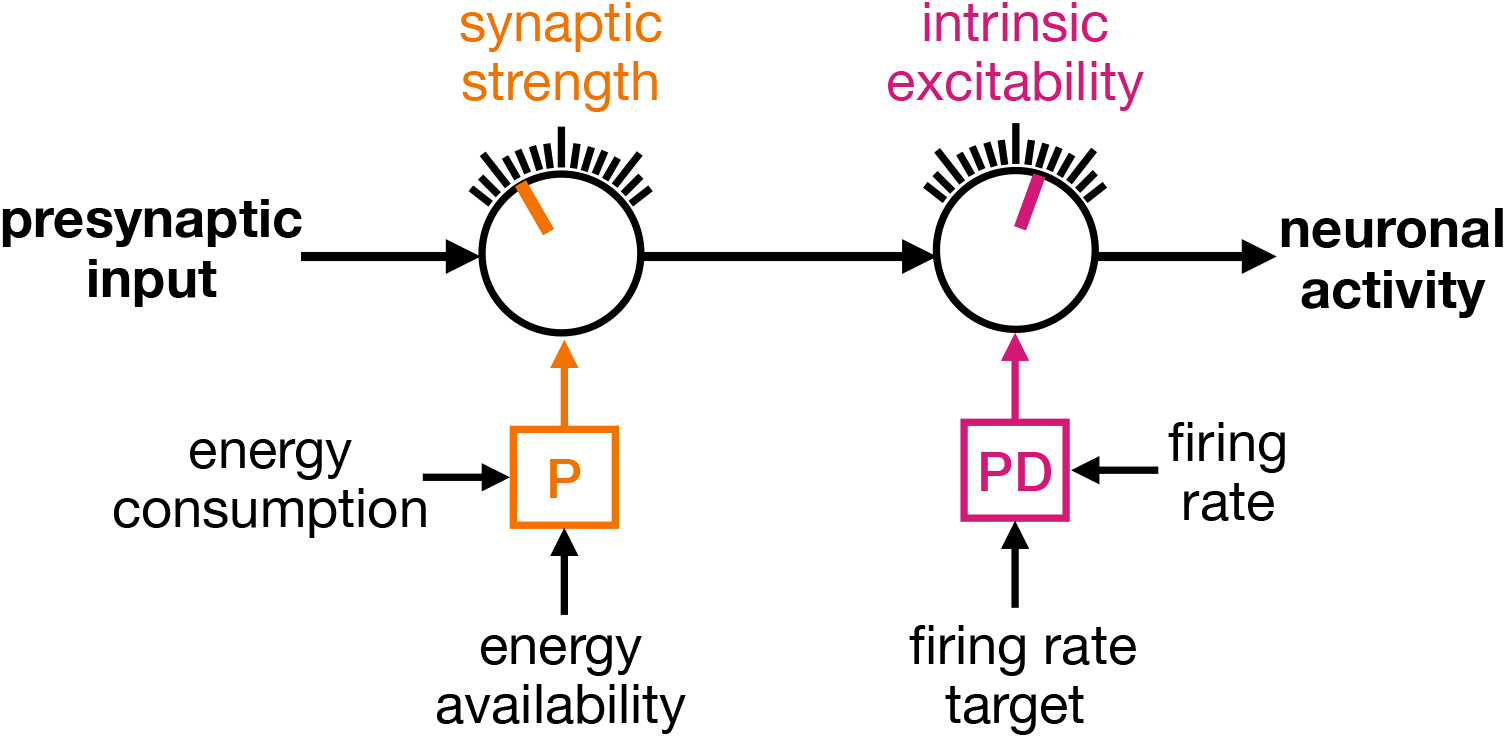
Energy-dependent control model of neuronal adaptation. Synaptic input is integrated by a postsynaptic neuron whose membrane and synaptic parameters determine its effective input–output gain. Energy scarcity reduces excitatory synaptic efficacy through energy-sensitive regulation, thereby lowering ATP demand. This change in synaptic efficacy is achieved by comparing energy consumption with available energy supply through a proportional controller. The resulting change in output activity is tracked by calcium-like activity signals, which drive homeostatic regulation of intrinsic excitability through a proportional–derivative controller. This feedback architecture allows the neuron to reduce energetic cost while maintaining firing within a functional range.

We test three implementations of this control architecture. First, in the *synapse-first* model, reduced energy availability directly downscales synaptic weights to lower ATP demand. This synaptic downscaling reduces output activity, which in turn triggers homeostatic increases in intrinsic excitability through higher membrane resistance and a depolarized resting potential. Thus, the model reduces energy demand while preserving firing rate. Second, in the *excitability-first* model, reduced energy availability acts first on intrinsic excitability. The resulting increase in firing rate then triggers compensatory synaptic down-scaling through activity-dependent homeostatic feedback. This implementation therefore tests whether an initially excitability-driven response can converge to the same energy-efficient regime.

Third, in the *variance-control* model, adaptation is constrained not only by energy availability and mean firing-rate homeostasis, but also by firing-rate variability (Triesch, 2004, 2005; Miller and Cannon, 2019). As in the *synapse-first* model, reduced energy balance downscales synaptic weights, whereas the resting potential is depolarized to stabilize mean firing rate. Membrane resistance, which controls synaptic integration and input–output variability, is regulated by the variability feedback signal. This third model therefore tests whether energy-efficient adaptation can also preserve higher-order features of neuronal output.

### Control-theoretic formulation of an energy-dependent integrate- and-fire neuron

To model how neurons could move toward energy-efficient parameter configurations, we translated the proposed *synapse-first* ordering into a dynamical control system. To do so, we extend an integrate-and-fire neuron, either LIF (Brunel and Van Rossum, 2007) or AdExp (Brette and Gerstner, 2005), with slow adaptation rules for synaptic weight scaling, resting membrane potential, and membrane resistance. Here, energy balance primarily regulates synaptic efficacy, whereas activity-dependent feedback controls intrinsic excitability. This separation allows the model to reduce energy demand while preserving output activity.

Synaptic efficacy is controlled by the slow energy-balance signal *ω_E_*(*t*). When ATP consumption exceeds available production, such that *ω_E_*(*t*) *< ω_E,_*_target_, the synaptic scaling factor *w*_scale_ decreases to reduce synaptic energy demand:

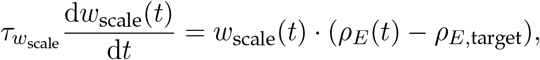

where *w*_scale_ scales excitatory and inhibitory synaptic conductances, *ε_w_*scale sets the time scale of energy-dependent synaptic adaptation, and *ω_E,_*_target_ = 1 denotes balanced ATP production and consumption. This rule corresponds to a simple proportional control law in which the change in the effector, d*w*_scale_(*t*)*/*d*t*, is proportional to the deviation of the target energy consumption and the actual energy consumption (magnitude of the energy deficit). We applied this common scaling factor to excitatory and inhibitory conductances, thereby lowering total synaptic drive while retaining the relative excitation–inhibition ratio. This is consistent with the tightly coupled excitation–inhibition balance observed in food restriction experiments (Padamsey et al., 2022, 2024). This energy-dependent synaptic adaptation is motivated by leptin-dependent AMPK–mTOR regulation of AMPAR-mediated synaptic efficacy during food restriction (Padamsey *et al*., 2024).

Reduced synaptic drive moves the neuron away from threshold and suppresses firing. Thus, a second control loop is required to stabilize neuronal output. In the *synapse-first* model, this stabilizing mechanism is a homeostatic increase in intrinsic excitability driven by an activity-control error:

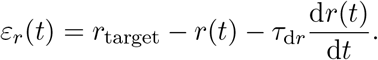

The proportional component measures the firing-rate deviation, whereas the derivative component opposes rapid changes and limits overshooting (Fig. S1). In control engineering terms, this formulation corresponds to a proportional-derivative (PD) controller, where the total change in the effector is the sum of a proportional term, *r*_target_ *→ r*(*t*), and a derivative term, d*r*(*t*)*/*d*t*. This proportional–derivative error regulates the two intrinsic variables found altered experimentally by food restriction, the resting potential *V*_rest_ and membrane resistance *R_m_* (Padamsey *et al*., 2022):

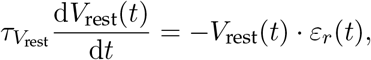

and:

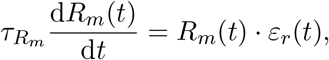

with distinct time constants *τ_V_rest__* and *ε_R_m__*, respectively. When activity falls below the target firing rate, *ε_r_*(*t*) *>* 0, the resting potential depolarizes and membrane resistance increases, thereby raising intrinsic excitability. An activity excess, *ε_r_*(*t*) *<* 0, reverses these adaptations. The resulting architecture assigns complementary objectives to the two control loops: metabolic feedback lowers the energetic cost of synaptic input, whereas intrinsic feedback restores output firing.

### Energy-dependent synaptic scaling and activity-dependent intrinsic adaptation reproduce observed food-restriction adaptations

We first test whether the *synapse-first* control architecture is able to reproduce the food-restriction phenotype of coordinated synaptic and intrinsic adaptations (Padamsey *et al*., 2022). In this model, reduced energy availability acts directly on synaptic strength, whereas deviations in output activity drive homeostatic changes in intrinsic excitability.

Under control conditions, energy supply matches the target energy budget (*s_E_* = 1), and the energy-balance signal remains close to *ω_E_* = 1 (Fig. 2A). Consequently, synaptic strength, firing rate, resting potential, and membrane resistance show no systematic drift (Fig. 2B–E). Thus, the control architecture preserves the initial neuronal operating point when ATP consumption matches production.

**Figure 2:**
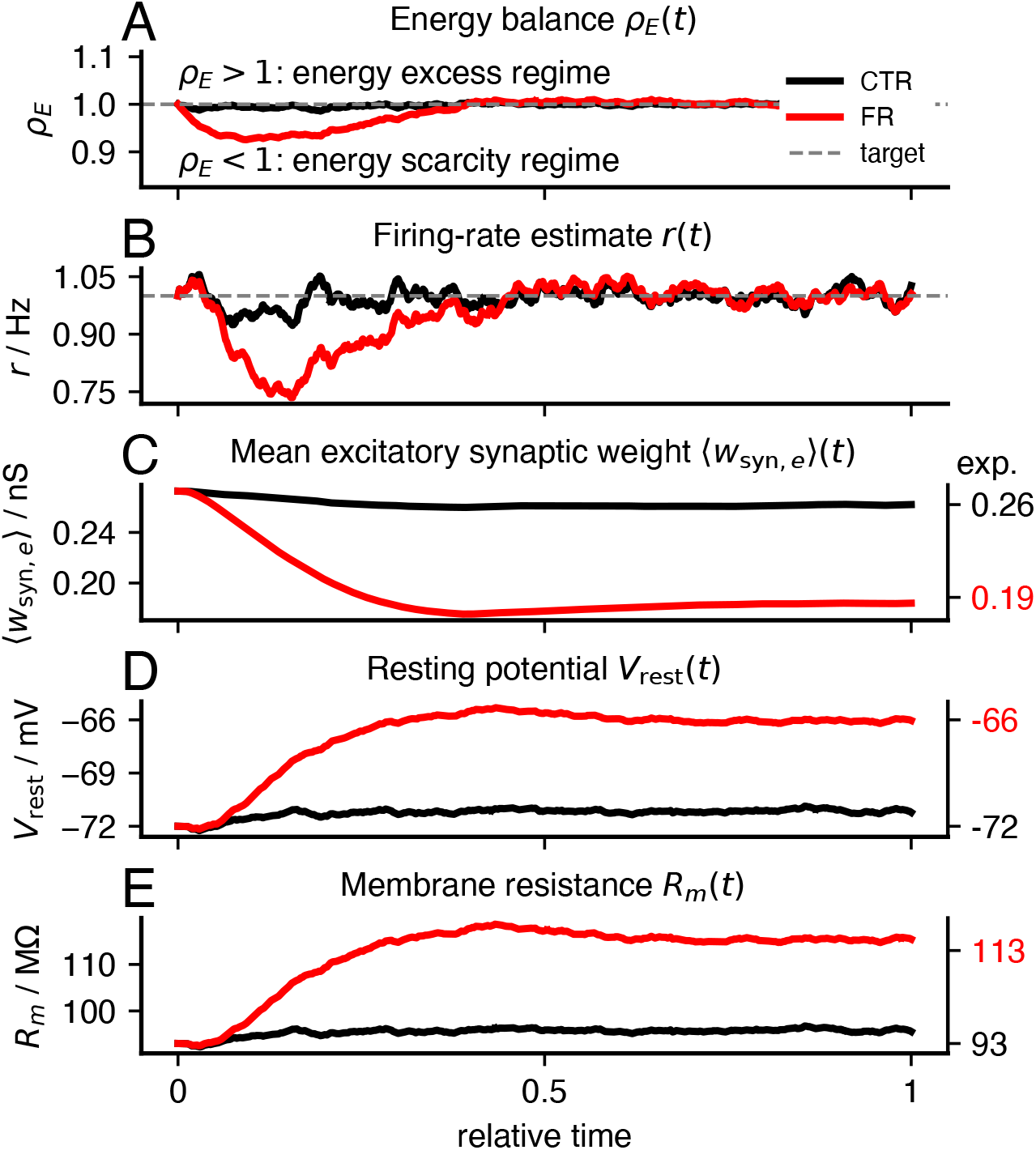
Energy-dependent synaptic scaling and activity-dependent intrinsic adaptation reproduce three parameter shifts experimentally observed after food-restriction. Simulation of the *synapse-first* control architecture under normal energy supply (control, CTR, black; *s_E_* = 1) and 15 % reduced energy supply (food restriction, FR, red; *s_E_*= 0.85). Experimental CTR and FR values are shown on the right side of panels C–E for comparison with simulated steady-state values. **A**, Energy-balance ratio *ω_E_*(*t*). In the CTR condition, energy balance remains close to *ω_E_* = 1, whereas reduced energy supply transiently drives the FR condition into an energy-deficient state. **B**, Firing rate. Synaptic down-scaling initially lowers firing rate in the FR condition before activity-dependent intrinsic adaptation restores firing toward the target rate. **C**, Mean excitatory synaptic efficacy. Energy-dependent synaptic scaling reduces synaptic drive in the FR condition until ATP consumption approaches the reduced supply. **D**, Resting potential. Homeostatic intrinsic adaptation depolarizes *V*_rest_ in the FR condition. **E**, Membrane resistance. Membrane resistance increases in parallel, raising intrinsic excitability and compensating for reduced synaptic drive.

Under food-restricted conditions, energy supply is reduced by 15 % (*s_E_* = 0.85) (Padamsey *et al*., 2022). This reduction in ATP supply causes an initial ATP deficit, driving the energy-balance signal below its target value. This control signal recruits synaptic downscaling, thereby lowering synaptic drive and ATP consumption (Fig. 2A,C). However, reduced synaptic drive transiently reduces firing below the target rate (Fig. 2B), thereby activating the intrinsic excitability control loop. Resting potential subsequently depolarizes and membrane resistance increases, compensating for the weaker input by raising intrinsic excitability (Fig. 2D,E). After this transient adaptation phase, the model eventually settles into a regime in which ATP demand matches the restricted supply and firing is largely recovered. When the relative adaptation time constants are constrained by the experimentally measured changes in *V*_rest_ and *R_m_* (see Supplements), the simulated steady state reproduces the direction and approximates the magnitude of the food-restriction phenotype (Padamsey *et al*., 2022). In total, this minimal *synapse-first* control architecture can move neurons toward the lower-energy operating regime observed experimentally (Padamsey *et al*., 2022) and can explain how reduced metabolic supply drives synaptic downscaling while intrinsic excitability compensates to preserve output activity.

Because the model separates synaptic and intrinsic control variables, it makes specific predictions. We therefore ask how adaptation would change if one intrinsic degree of freedom were constrained; for example by blocking resting-potential adaptation. In this case, energy-dependent synaptic scaling still reduces synaptic drive under food restriction (Fig. S2A–C). However, the loss of resting-potential compensation shifts homeostatic adaptation onto the remaining intrinsic degree of freedom. Membrane resistance increases more strongly than in the unconstrained model, reaching approximately 170 MΛ instead of 113 MΛ when both intrinsic mechanisms are available (Fig. S2E). Thus, blocking one compensatory pathway forces the model to restore firing through the remaining intrinsic degree of freedom. This perturbation provides a direct experimental prediction. If resting-potential depolarization is prevented during reduced energy availability, for example through manipulating the extracellular ionic balance, neurons should show an exaggerated compensatory increase in membrane resistance. More generally, the model predicts that food-restriction adaptations are not independent parameter changes but coupled compensatory movements within a constrained energy–activity control system.

### *Excitability-first* control produces transient hyperactivity and reduces coding efficiency

The *synapse-first* architecture assumes that energy balance primarily regulates synaptic efficacy, whereas intrinsic excitability stabilizes output firing. This causal ordering is motivated by leptin-dependent AMPK–mTOR regulation of AMPAR-mediated synaptic efficacy during food restriction (Padamsey et al., 2022, 2024). However, the available experiments do not establish a temporal precedence. We therefore consider what would happen if the mapping of the two feedback signals were reversed: in an alternative *excitability-first* architecture energy balance controls intrinsic excitability, potentially through leak, HCN, or voltage-gated conductances, and firing-rate feedback then adjusts synaptic strength

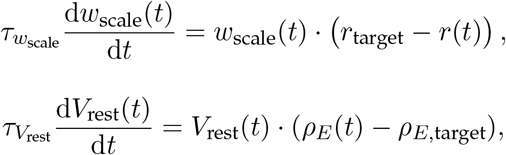

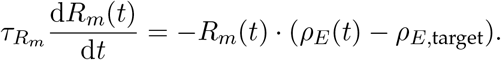

Under reduced energy supply, the metabolic controller first increases intrinsic excitability, causing firing to overshoot before activity-dependent synaptic downscaling restores the target firing rate (Fig. 3). The two architectures therefore predict opposite early responses to energy scarcity: transient hyperactivity in the *excitability-first* model and transient firing suppression in the *synapse-first* model.

**Figure 3:**
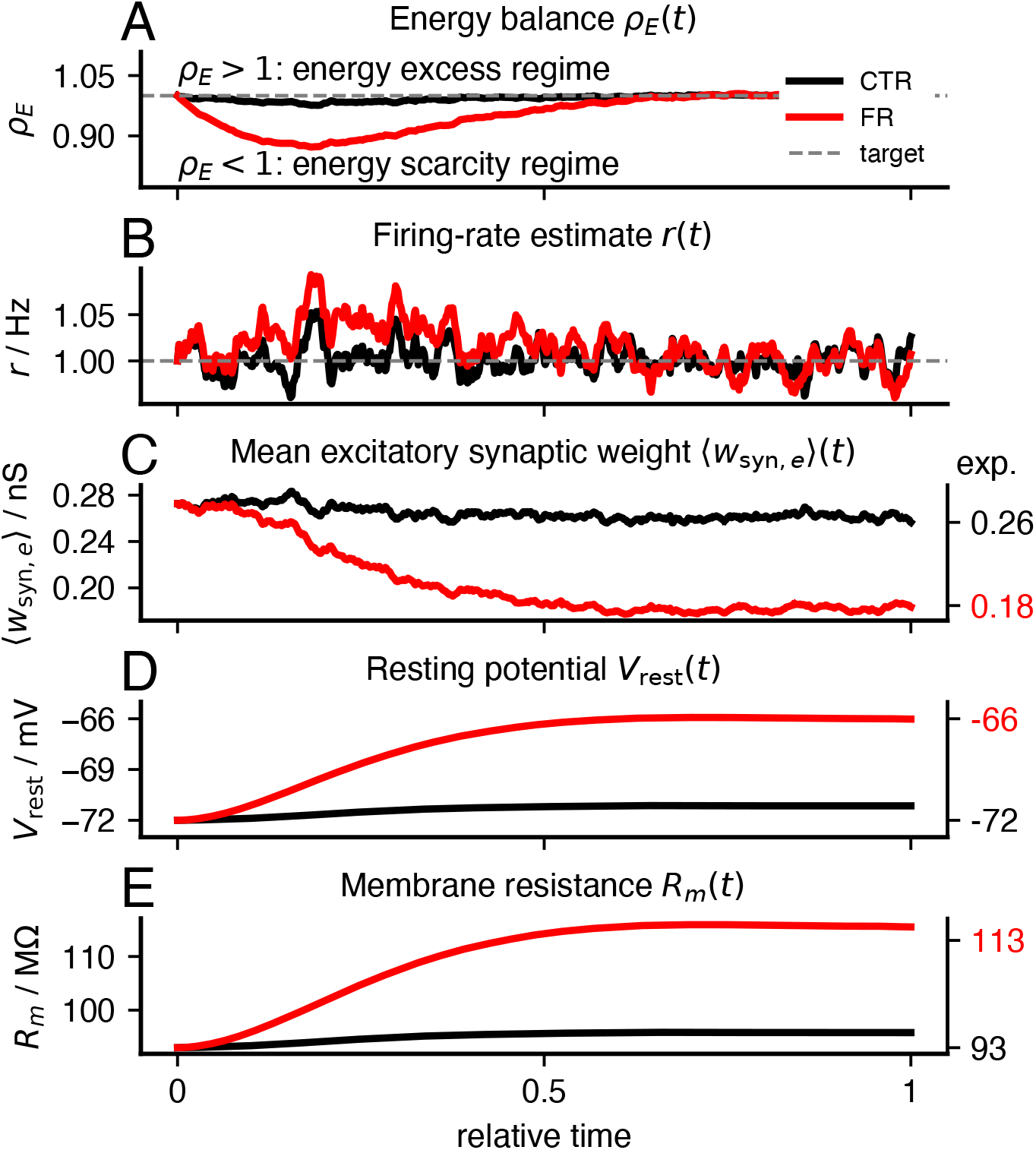
*Excitability-first* control produces transient hyperactivity. Simulation of the *excitability-first* control architecture under normal energy supply (control, CTR, black; *s_E_* = 1) and 15 % reduced energy supply (food restriction, FR, red; *s_E_* = 0.85). In this formulation, energy balance directly regulates intrinsic excitability, whereas activity-dependent feedback regulates synaptic strength. **A**, Energy-balance signal *ω_E_*(*t*). In the FR condition, restricted supply displaces the energy-balance signal from its target. **B**, Firing-rate estimate *r*(*t*). Unlike the *synapse-first* model, the *excitability-first* architecture produces a transient increase in firing before activity-dependent synaptic compensation restores firing toward the target rate. **C**, Mean excitatory synaptic weight. Firing-rate feedback subsequently reduces synaptic strength and returns activity toward its target value. **D– E**, Resting potential and membrane resistance. Energy-dependent intrinsic adaptation changes *V*_rest_ and *R_m_* under reduced energy supply, increasing excitability before synaptic compensation stabilizes activity.

This transient increase in firing rate in the *excitability-first* model increases energy consumption relative to the *synapse-first* model (Fig. 4A). We ask whether this higher energetic cost is accompanied by improved coding performance. To test this, we evaluate both trajectories in the oriented-grating discrimination task used in the food-restriction experiments (Padamsey *et al*., 2022) and modeled previously (Sommer *et al*., 2026). The *synapse-first* trajectory maintains higher orientation selectivity at comparable energy consumption, whereas the *excitability-first* trajectory transiently occupies a lower-selectivity regime (Fig. 4B). A plausible explanation is that *synapse-first* control initially reduces synaptic drive and associated background fluctuations while intrinsic excitability remains unchanged. In contrast, *excitability-first* control increases sensitivity to noise-driven spiking before synaptic downscaling occurs, potentially reducing stimulus specificity and orientation selectivity (Sommer *et al*., 2026). Thus, the additional energy consumed during *excitability-first* adaptation does not translate into improved sensory selectivity. Instead, the *synapse-first* trajectory provides a more favorable path through the energy–information trade-off, reducing energy demand while preserving orientation selectivity over a broader range of the adaptation trajectory.

**Figure 4:**
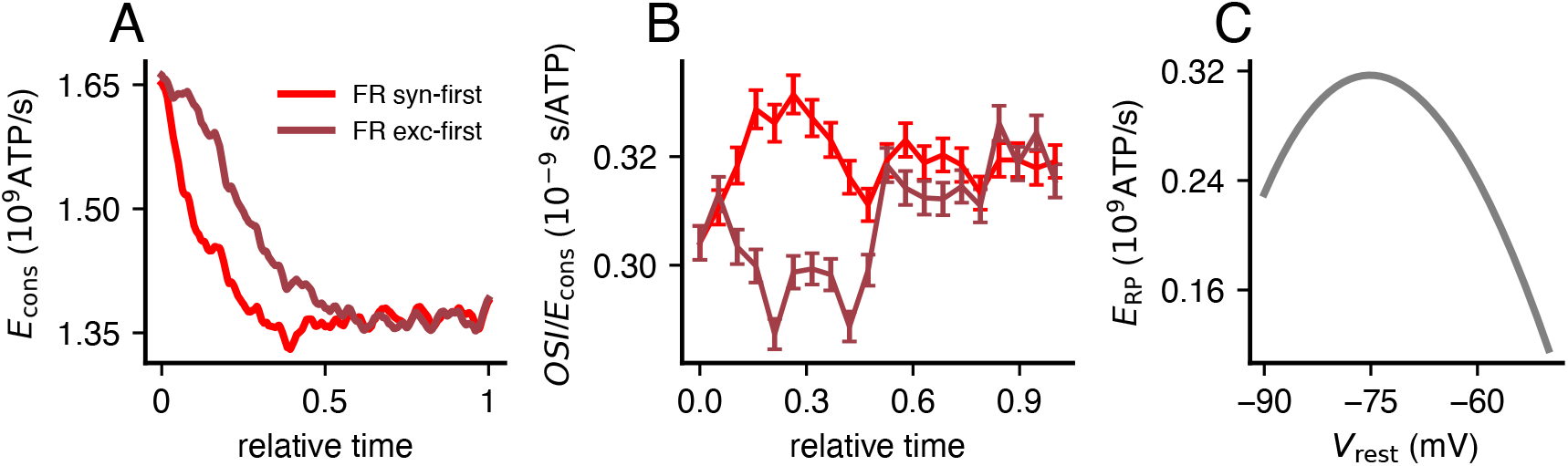
*Synapse-first* adaptation follows a more favorable energy–information trajectory than *excitability-first* adaptation. Comparison of the *synapse-first* and *excitability-first* control architectures along the food-restriction adaptation trajectory. **A**, Total energy consumption over relative adaptation time. The *excitability-first* model shows higher transient energy consumption and reaches its steady state more slowly than the *synapse-first* model. **B**, Orientation selectivity per energy consumption (*OSI/E*_cons_) during an oriented-grating discrimination task (Padamsey *et al*., 2022; Sommer *et al*., 2026). Error bars indicate the standard error of the mean (SEM) across 2000 simulations. The *synapse-first* model achieves higher orientation selectivity per energy consumption over a broader portion of the trajectory than the *excitability-first* model. **C**, Resting-potential energy demand as a function of *V*_rest_. The non-monotonic dependence of *E*_RP_ on *V*_rest_ illustrates why direct energy-based control of intrinsic excitability can produce counterintuitive adaptation dynamics.

### Variance-based intrinsic control introduces instability without clear experimental support

To assess whether food restriction alters spike-train variability, we analyzed *in vivo* voltage recordings obtained during natural-scene stimulation (Padamsey *et al*., 2022). The dataset comprised neuronal voltage-trace recordings from 40 control (CTR) and 37 food-restricted (FR) cells. We identified spikes using a peak prominence of 42 and a minimum separation of 10 ms; leaving 26 CTR and 17 FR cells with sufficient spikes for interspike-interval (ISI) analyses.

Mean firing rates during natural scene stimulation were low and did not differ significantly between CTR and FR neurons (0.18 versus 0.12 Hz; Mann–Whitney U test, *p* = 0.28; Fig. 5A). ISI variance was numerically lower under food restriction, although the difference did not reach statistical significance (37.2 versus 18.3 s^2^; Mann–Whitney U test, *p* = 0.068; Fig. 5B). Because absolute ISI variance depends on firing rate, we additionally quantified rate-normalized irregularity using *CV*_ISI_. The resulting values were nearly identical between conditions (1.12 for CTR and 1.11 for FR; Welch’s *t*-test, *p* = 0.96; Fig. 5C). Therefore, neither absolute nor rate-normalized ISI variability changed significantly under food restriction. Although the numerical reduction in ISI variance is compatible with altered spike-train statistics, the present experimental data cannot determine whether spike-train variability is directly regulated as a homeostatic target.

**Figure 5:**
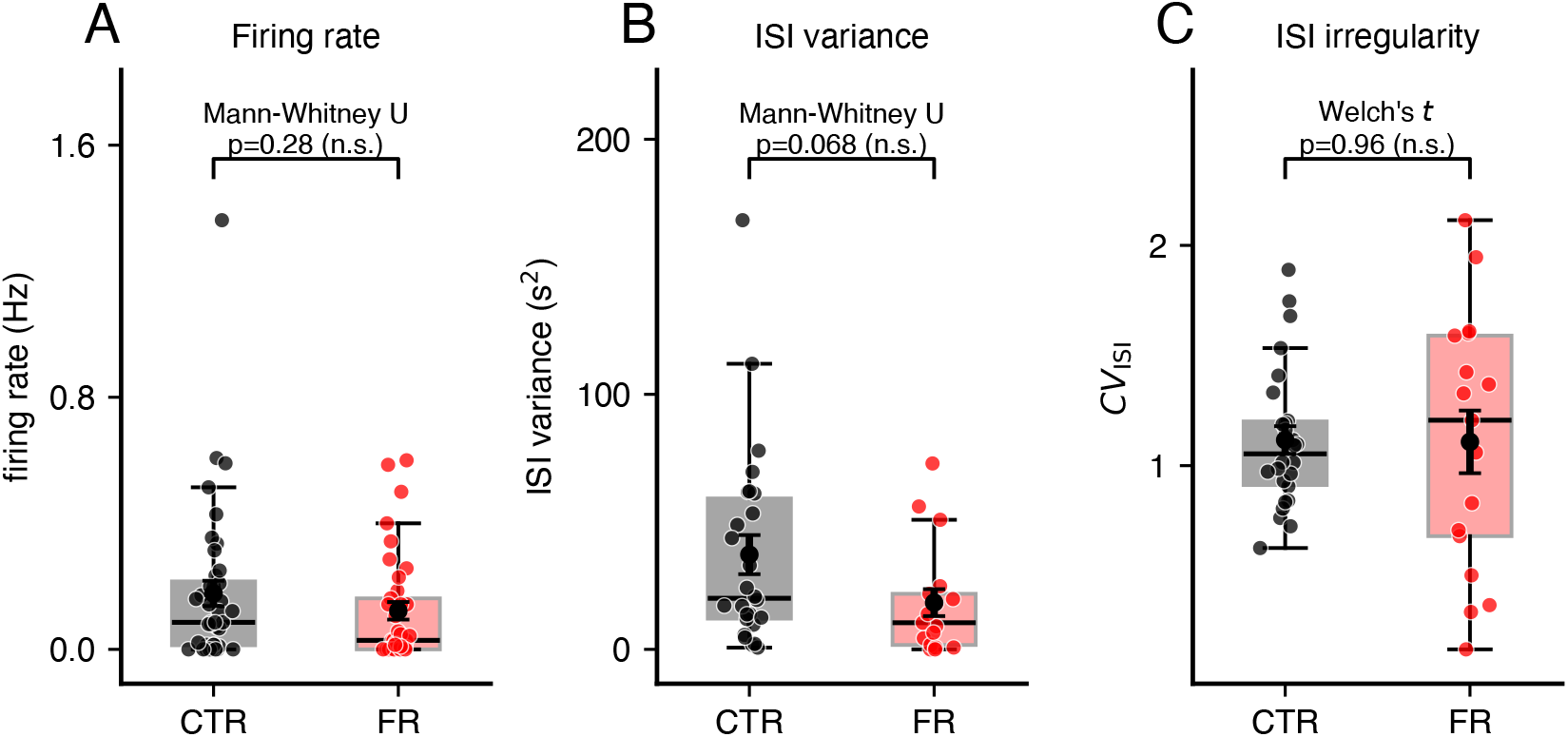
Firing rate and spike-train variability during natural scene stimulation. **A**, Mean firing rates of neurons from control (CTR, black) and food-restricted (FR, red) animals during natural scene stimulation. Individual points indicate single cells; box plots show the median and interquartile range. Mean firing rates were low in both groups and did not differ significantly. **B**, Interspike-interval (ISI) variance for CTR and FR neurons. Although ISI variance was numerically lower in FR neurons, the difference was not statistically significant. **C**, Coefficient of variation of interspike intervals (*CV*_ISI_), providing a rate-normalized measure of spike-train irregularity. *CV*_ISI_ did not differ between CTR and FR neurons. Together, these results provide no evidence for a systematic change in firing-rate variability or spike-train irregularity under food restriction during natural scene stimulation.

In the AdExp simulations, food-restriction parameter combinations produced slightly more regular spike trains than the control configuration. The coefficient of variation of interspike intervals decreased from *CV*_ISI_ = 0.68 in the CTR setting to *CV*_ISI_ = 0.59 in the FR setting. (For comparison, the standard leaky integrate-and-fire (LIF) model yielded *CV*_ISI_ = 0.99.) Because the mean firing rate was constrained by the homeostatic controller, this reduction mainly reflected a lower ISI variance, as observed in the experimental data (Fig. 5B), rather than a change in average activity. However, the large variability in the experimental *CV*_ISI_ estimates (Fig. 5C) makes it difficult to determine whether such variability is actively controlled, and if so, how strongly.

Although the experimental data do not establish variability as an additional homeostatic target, we test whether assigning distinct higher-order activity statistics to intrinsic parameters could improve control of neuronal output. Because input resistance shapes synaptic integration through the membrane time constant and may therefore influence spike-train variability in addition to the mean firing rate (Triesch, 2007; Miller and Cannon, 2019), we extend the *synapse-first* architecture and couple *V*_rest_ to mean-rate feedback and *R_m_* to firing-rate variance, while retaining metabolic control of synaptic strength:

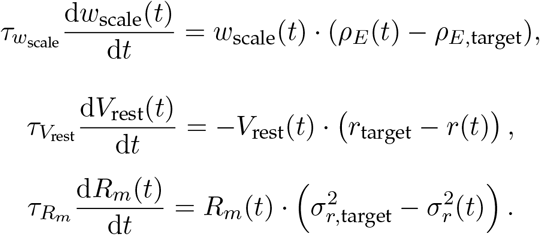

In principle, this architecture could enable a neuron to regulate multiple output features simultaneously, including mean activity and spiking variability. However, separating the control targets did not separate their physiological effects in our simulations. The model failed to converge robustly likely because changes in *R_m_* altered both variability and mean excitability, thereby perturbing the firing rate regulated through *V*_rest_ (Supplementary Fig. 6). We conclude that this instability likely arises because resting potential and membrane resistance do not affect independent output dimensions, thereby producing interacting feedback loops that could oppose one another. Similar instability is observed when the squared coefficient of variation of firing rate, 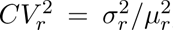, is used as the controlled variability measure instead of firing-rate variance. Thus, the instability is not specific to the absolute scale of firing-rate variance, but reflects the broader difficulty of independently controlling mean activity and variability through coupled intrinsic parameters

Together, these findings suggest that explicitly controlling variability is not necessary to reproduce the food-restriction-like adaptations and may introduce additional instability. Combined with the ambiguous experimental support for variability control the results therefore do not rule out variance regulation, but they provide insufficient evidence to include it as a primary control objective in the main model.

## Discussion

Neurons must continuously balance several competing demands: they need to remain responsive to input, preserve information-processing capacity, stabilize activity, and keep energy consumption within available metabolic limits. These objectives can be regulated through many cellular and circuit-level “knobs”, including synaptic efficacy, dendritic integration, intrinsic properties such as spike threshold, resting potential and membrane resistance, and, at the network level, excitation–inhibition balance (Turrigiano *et al*., 1998; Desai *et al*., 1999; O’Leary and Wyllie, 2011; Debanne *et al*., 2019; Yang and Prescott, 2023). This makes neuronal homeostasis a highly degenerate control problem, in which different parameter combinations may achieve similar functions or firing rates while differing in energy demand and coding performance (Sommer *et al*., 2026). Here, we focused on a restricted but experimentally motivated subset of this regulatory landscape: the coordination of synaptic efficacy, resting potential, and membrane resistance under reduced energy availability (Padamsey *et al*., 2022).

**Figure 6:**
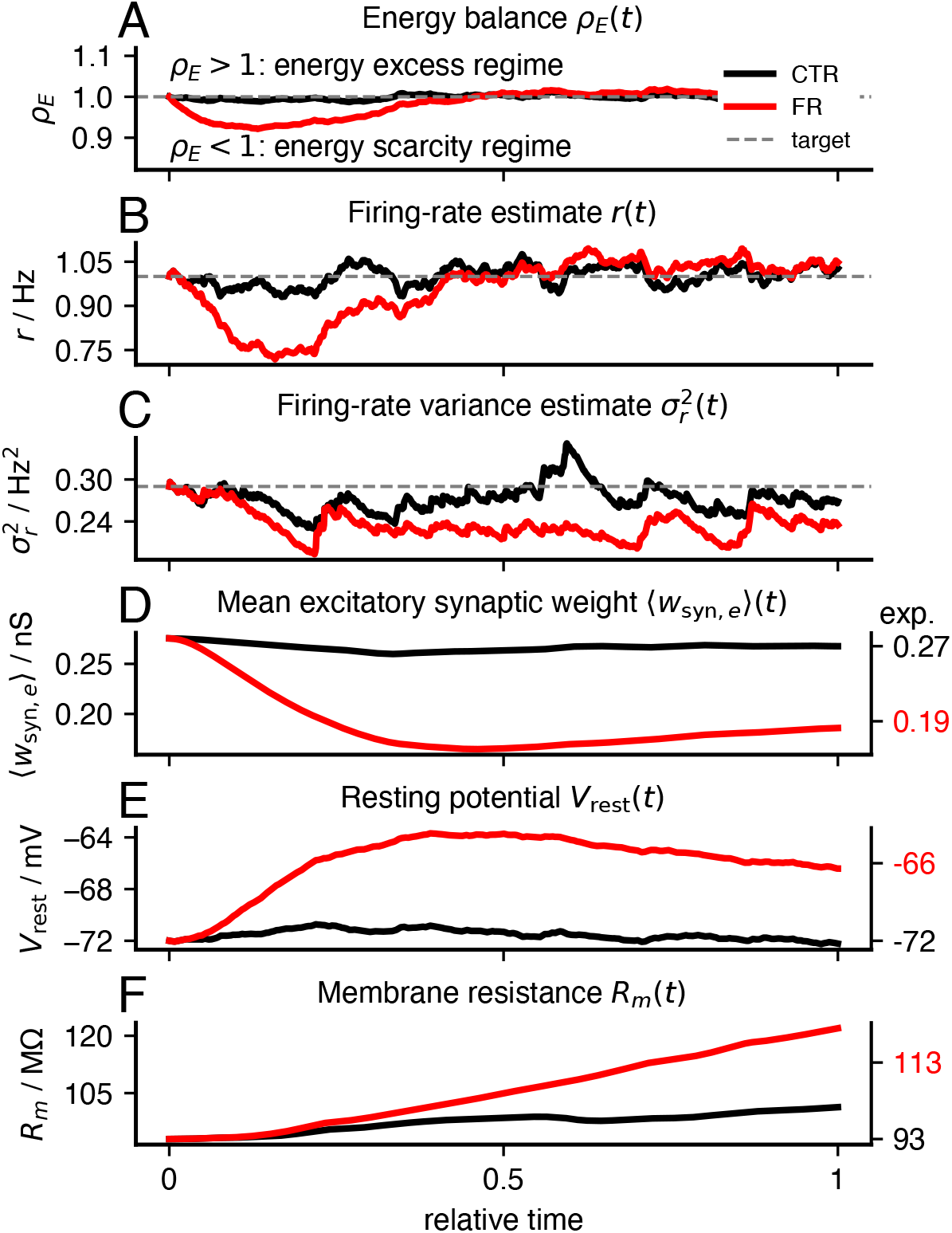
Variance-based intrinsic control does not converge robustly. Simulation of an alternative control architecture in which energy balance regulates synaptic scaling, mean firing-rate feedback regulates *V*_rest_, and firing-rate variance feedback regulates *R_m_*. **A**, Energy-balance signal *ω_E_*(*t*) under normal energy supply (control, CTR, black; *s_E_* = 1) and reduced energy supply (food restriction, FR, red; *s_E_* = 0.85). **B**, Low-pass filtered firing-rate estimate *r*(*t*). **C**, Firing-rate variance estimate 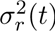 relative to the target variance 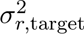. **D**, Mean excitatory synaptic weight *w*_syn_*_,e_*(*t*), which is reduced under food restriction by energy-dependent synaptic scaling. **E**, Resting potential *V*_rest_(*t*), regulated by mean firing-rate feedback. **F**, Membrane resistance *R_m_*(*t*), regulated by variance feedback. Because *V*_rest_ and *R_m_* both affect mean excitability and spiking variability, their interacting adaptation produces unstable or non-convergent dynamics. This indicates that variance-based intrinsic control is less robust than the firing-rate-homeostatic architecture used in the *synapse-first* model.

### Coordinating synaptic and intrinsic plasticity under energy scarcity

Food restriction induces coordinated weakening in synaptic input and increased intrinsic excitability in cortical neurons (Padamsey *et al*., 2022, 2024). This coordinated response is consistent with a reduction in ATP-expensive synaptic demand and compensatory changes in membrane resistance and resting potential that preserve output activity. However, measurements obtained after adaptation cannot determine which change initiated the transition. Among the many possible ways in which neurons could adjust their properties, we considered two simplified limiting cases that coordinate energy demand with firing-rate homeostasis in opposite directions: metabolic stress may first suppress synaptic efficacy and recruit intrinsic excitability compensation, or it may first alter intrinsic excitability and subsequently recruit homeostatic synaptic scaling. Because these plasticity classes engage partly distinct pathways and need not serve the same functional objective (Wu *et al*., 2020; Wen *et al*., 2025), their ordering has consequences for both transient activity and energetic cost. We additionally considered whether membrane resistance could regulate spike-train variability rather than mean firing. Although all three architectures, in principle, can approach related steady states, their adaptation trajectories provide distinct mechanistic predictions. We therefore compared three simplified architectures: a *synapse-first* model, in which metabolic state regulates synaptic efficacy and activity feedback regulates intrinsic excitability; an *excitability-first* model, in which metabolic state regulates intrinsic excitability and activity feedback regulates synaptic strength; and a *variance-control* model, in which membrane resistance is regulated by spiking variability rather than mean firing rate.

### Experimental evidence supports a *synapse-first* interpretation

AMPAR-mediated excitatory synaptic efficacy ((Wang et al., 2006; Potter et al., 2010; Ahmadi and Roy, 2016; Herzig and Shaw, 2018; Gonźalez et al., 2020). The resulting reduction in. glutamatergic drive would directly lower one of the major activity-dependent components of neuronal ATP consumption.

The available experiments do not, however, uniquely resolve temporal order. Metabolic signaling pathways can also influence intrinsic excitability, and leptin can modulate ion channels that affect resting potential, membrane resistance, and dendritic integration, including leak, HCN, and voltage-gated conductances (Harvey, 2007; Wahl-Schott and Biel, 2008). An *excitability-first* scenario therefore remains biologically possible. In such a model, metabolic stress would first alter excitability, and activity-dependent synaptic scaling would subsequently restore firing rates (Turrigiano *et al*., 1998). However, this ordering produces a transient phase of hyperexcitability (Fig. 3B). Such transient hyperactivity is suboptimal because it increases neuronal energy demand and is accompanied by reduced orientation selectivity at comparable energy consumption (Fig. 4A–B).

Experimental support for variance-based homeostatic control remains ambiguous. During natural scene stimulation, *in vivo* recordings did not reveal a reliable food-restriction-induced change in spike-train variability or interspike-interval irregularity. Although variability can influence information transmission, these data are not sufficient to determine whether firing-rate variance is actively regulated as a primary homeostatic target in this paradigm.

### Simulation evidence favors *synapse-first* control

The simulation results further support the *synapse-first* architecture as the most parsimonious control strategy by distinguishing the adaptation trajectories rather than only their endpoints. Assigning metabolic feedback to synaptic efficacy immediately targets a major endpoints. Assigning metabolic feedback to synaptic efficacy immediately targets a major shortage, whereas intrinsic feedback subsequently restores firing by increasing membrane resistance and depolarizing the resting potential. Furthermore, this ordering reduces energy demand during adaptation because it prevents an ATP-costly firing-rate overshoot produced by *excitability-first* control. At the same time, the *synapse-first* trajectory maintains higher orientation selectivity in discrimination tasks than the *excitability-first* trajectory for the same amount of ATP consumption over a broad range of the adaptation process (Fig. 4A–B). The simulations therefore favor *synapse-first* control because it reaches the food-restriction-like state through a more economical transient regime.

The energetic leverage of the two mechanisms also differs. In the energy budget, synaptic transmission contributes more strongly to activity-dependent ATP consumption than resting-potential maintenance. Directly scaling excitatory synaptic input therefore provides a more effective first response to reduced energy availability. By contrast, *excitability-first* control must rely on subsequent activity-dependent synaptic scaling to substantially reduce total energy consumption, making the architecture less direct and more prone to transient overshoot in firing rate and energy consumption.

In addition, metabolic control of the *excitability-first* architecture is less well specified because the cost of maintaining the resting potential *E*_RP_ depends non-monotonically on *V*_rest_ and *R_m_* (Fig. 4C). Depending on the initial operating point, either depolarization or hyperpolarization might reduce this cost. The *excitability-first* architecture therefore acts through a more ambiguous energetic signal and requires delayed synaptic compensation, providing a plausible explanation for its firing-rate overshoot and lower dynamical robustness.

The *variance-control* model provided a possible extension in which membrane resistance regulates spiking variability, a feature potentially relevant for information transmission (Triesch, 2004, 2005, 2007). However, this architecture did not converge robustly in our simulations, likely because *R_m_*and *V*_rest_ do not control fully independent output dimensions: both intrinsic parameters affect mean activity and variability, so correction of one statistic can perturb the other and generate opposing feedback. Within this reduced model, *R_m_* cannot be treated as a dedicated actuator of firing variability. In general, however, *R_m_*controls the integration of synaptic inputs in time and so influences the capacity of a neuron to perform temporal coding; higher *R_m_* makes a cell less sensitive to coincident inputs. In V1 this appears not to be a major factor (Sommer *et al*., 2026), but might allow for a different control approach in neurons with different coding tasks.

Together, the molecular and computational evidence identifies *synapse-first* control as the most parsimonious explanation of food-restriction adaptation. It assigns the primary reduction in energetic cost to synaptic scaling and the stabilization of neuronal output to intrinsic plasticity. This division should not be interpreted as an exclusive molecular pathway: AMPK, mTOR, calcium-dependent signaling, ion-channel regulation, and receptor trafficking are likely to interact across both forms of adaptation. Rather, the *synapse-first* model presents a minimal control principle that captures the dominant direction of adaptation under restricted energy supply: Neurons can economize signaling through synaptic downscaling while using intrinsic plasticity to retain functional activity.

### Experimental tests to distinguish *synapse-first* from *excitability-first* control

The competing models are distinguished primarily by their temporal dynamics rather than their final parameter values. Resolving their causal ordering will therefore require repeated measurements of synaptic efficacy, membrane properties, firing, and spike-train variability as metabolic stress develops, combined with selective perturbations of the corresponding plasticity pathways.

Under *synapse-first* control, excitatory synaptic weakening should precede the increase in intrinsic excitability. Preventing AMPAR downscaling through manipulation of leptin– AMPK–mTOR signaling or receptor trafficking should therefore attenuate the loss of coding precision, but also maintain synaptic ATP demand and limit energy savings. By contrast, preventing the subsequent intrinsic compensation should leave synaptic weakening intact and expose a sustained reduction in firing and sensory performance.

Under *excitability-first* control, changes in membrane resistance or resting potential should instead emerge before synaptic weakening and produce a transient firing-rate overshoot. Blocking the later synaptic response should convert this transient into persistent hyperactivity accompanied by elevated ATP demand. Early firing dynamics therefore provide a key experimental discriminator: initial suppression supports *synapse-first* control, whereas initial overshooting supports an *excitability-first* sequence.

A decisive experiment should therefore combine temporally resolved physiological measurements with pathway-specific perturbations from the onset of food restriction. Relating the resulting adaptation sequence to ATP demand and coding precision would reveal whether tuning degradation reflects an actively regulated energy–information trade-off or a nonspecific failure caused by metabolic stress.

### Bidirectional energy-dependent regulation of synaptic strength

Our energy-dependent framework may also explain homeostatic adaptations following prolonged neuronal silencing. Experiments using tetrodotoxin (TTX) silencing action potential generation and transmission have demonstrated increased excitatory synaptic strength and membrane resistance following activity deprivation (Turrigiano *et al*., 1998; Desai *et al*., 1999; Wen *et al*., 2025). In our model, reduced neuronal activity decreases ATP consumption, creating a relative energy surplus that promotes synaptic upscaling, while activity-dependent feedback increases membrane resistance. An exploratory simulation reproduced these qualitative adaptations (Supplementary Fig. S3), suggesting that the same feedback mechanisms may operate under both energy scarcity and activity deprivation.

This interpretation is consistent with the hypothesis of Chintaluri and Vogels (2023) who proposed that neurons may increase metabolically costly spiking during periods of low energy demand to maintain metabolic homeostasis and limit oxidative stress. Our framework suggests a complementary mechanism in which synaptic downscaling reduces communication costs under energy scarcity, whereas synaptic upscaling increases the potential energetic cost of communication under conditions of relative energy abundance. Whether metabolic feedback directly contributes to TTX-induced homeostatic plasticity remains to be established.

### Disrupted energy regulation promotes pathological excitability

Neuronal energy supply and demand have received increasing attention because metabolic dysfunction is closely linked to neurological disease (Bazzigaluppi *et al*., 2017). The same adaptations that stabilize neuronal function during moderate energy scarcity may become insufficient or maladaptive when metabolic stress exceeds the compensatory range. Neurons are particularly susceptible because their high ATP demand is supported by limited energetic reserves (Klemmensen *et al*., 2024). When energetic stress is transient, neurons can compensate by adjusting synaptic and intrinsic parameters. When it becomes chronic, however, impaired mitochondrial function can impair ion homeostasis, alter excitability, and contribute to progressive neuronal dysfunction (Klemmensen *et al*., 2024; Kö hler-Soĺıs and Schirmeier, 2025). Such long-term energy deficits could shift neuronal operating points toward chronically depolarized membrane potentials, altered reversal potentials, or disrupted excitation–inhibition balance, thereby increasing network excitability and seizure susceptibility (Le Masson *et al*., 2014). Such mechanisms may be relevant for disorders in which metabolic dysfunction and excitability changes interact, including epilepsy and neurodegenerative diseases such as Alzheimer’s disease (Engl and Attwell, 2015). In this view, disease emerges from a failure of the control mechanisms that normally balance metabolic demand against stable information processing.

Integrating the energy-dependent integrate-and-fire model into recurrent networks would test whether chronic, fluctuating, or spatially heterogeneous energy deficits promote bursting and destabilize population activity. This extension could also distinguish the consequences of persistently low supply from rapid metabolic fluctuations, a distinction that may be relevant to the anticonvulsant effects of interventions such as ketogenic diets (Ułamek-Kozioł *et al*., 2019). More broadly, linking cellular energy control to network dynamics may therefore reveal when homeostatic adaptation ceases to protect information processing and instead promotes pathological excitability.

## Conclusion

Together, our results suggest that cortical neurons adapt to energy scarcity through a coordinated control strategy that separates energetic and functional objectives. Reduced metabolic supply primarily downscales ATP-expensive excitatory synaptic input, whereas intrinsic plasticity compensates for the resulting loss of drive to preserve firing within a functional range. This *synapse-first* architecture provides a simple mechanistic explanation for how food restriction can reduce energetic cost while degrading coding precision. More broadly, these findings support the view that neuronal parameters are not optimized for information transmission alone, but are constrained by a dynamic trade-off between energy consumption, coding performance, and functional stability.

**Table 1:** Parameters for the energy-dependent LIF and AdExp neuron models and homeostatic control mechanisms.

| Parameter | Value | Unit | Reference |
| --- | --- | --- | --- |
| **Neuron model parameters** |  |  |  |
| $C_m$ | 100 | pF | Padamsey *et al.* (2022) |
| $V_{\text{thresh}}$ | -50 | mV | Zerlaut and Destexhe (2017) |
| $V_{\text{reset}}$ | $V_{\text{rest}}$ | mV | |
| $\Delta_T$ | 2 | mV | Zerlaut and Destexhe (2017) |
| $V_T$ | $V_{\text{thresh}} - 2.5 \cdot \Delta_T$ | mV | Zerlaut and Destexhe (2017) |
| $\tau_{\text{ad}}$ | 500 | ms | Zerlaut and Destexhe (2017) |
| $a_{\text{ad}}$ | 4 | nS | Zerlaut and Destexhe (2017) |
| $b_{\text{ad}}$ | 0.02 | nA | Zerlaut and Destexhe (2017) |
| **Activity trace and control parameters** |  |  |  |
| $\tau_{\text{Ca}^{2+}}$ | 1 000 | ms | |
| $A_{\text{Ca}^{2+}}$ | 1.0 | Hz | |
| $r_{\text{target}}$ | 1.0 | Hz | Padamsey *et al.* (2022) |
| $\rho_{E,\text{target}}$ | 1.0 | – | |
| $s_E$ | CTR: 1.0, FR: 0.85 | – | Padamsey *et al.* (2022) |
| $E_{\text{target,ATP}}$ | $1.6 \cdot 10^9$ | ATP/s | |
| $\tau_r$ | 10 000 | ms | |
| $\tau_{\text{dr}}$ | 2 500 | ms | |
| $\tau_E$ | 10 000 | ms | |
| $\tau_{V_{\text{rest}}}$ | 50 000 | ms | |
| $\tau_{R_m}$ | 20 000 | ms | |
| $\tau_{w_{\text{scale}}}$ | 4 000 | ms | |
| $\sigma_{r,\text{target}}^2$ | 0.29 | Hz<sup>2</sup> | |

**Table 2:**
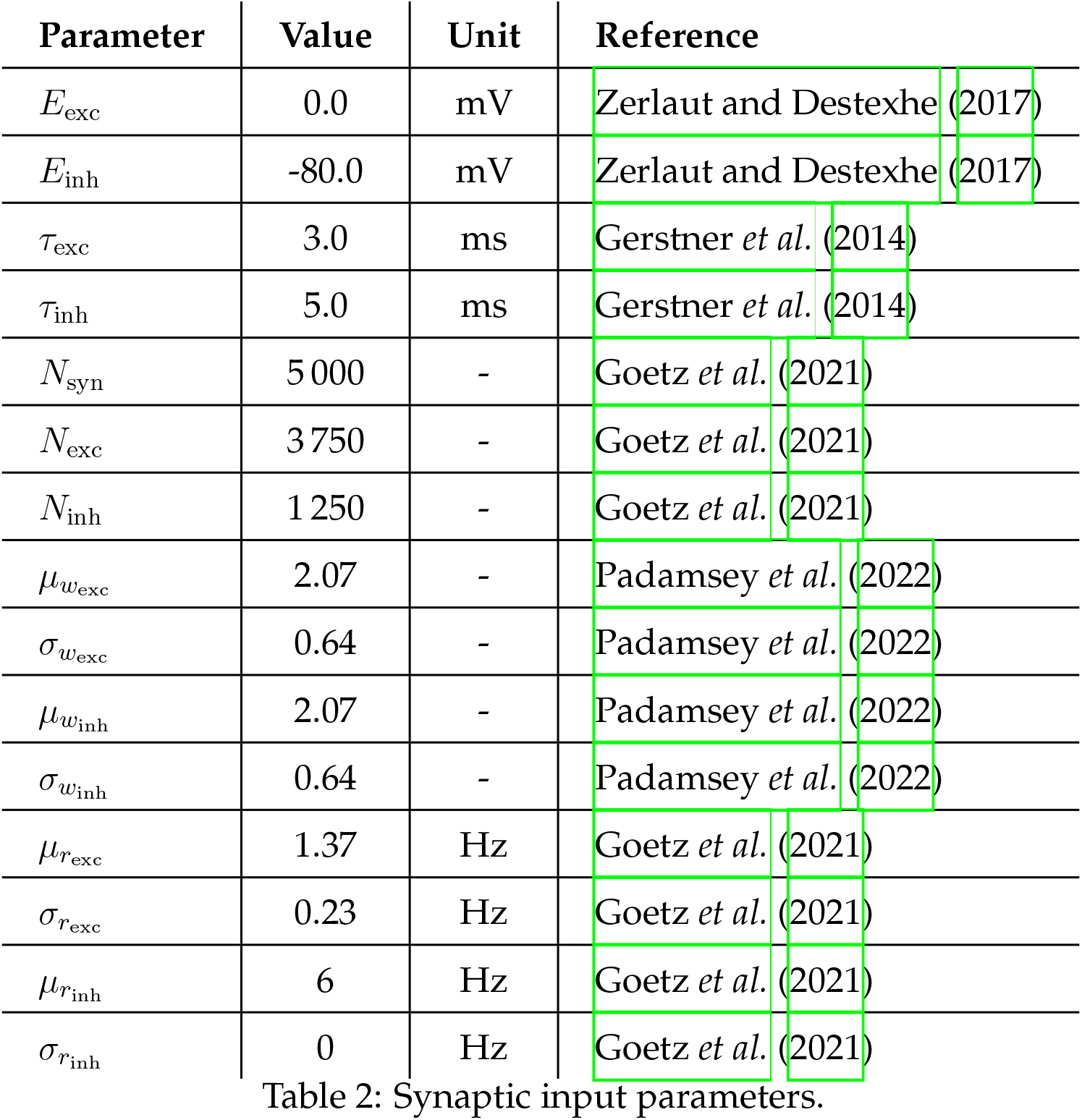
Synaptic input parameters.

| Parameter | Value | Unit | Reference |
| --- | --- | --- | --- |
| $E_{\text{exc}}$ | 0.0 | mV | Zerlaut and Destexhe (2017) |
| $E_{\text{inh}}$ | -80.0 | mV | Zerlaut and Destexhe (2017) |
| $\tau_{\text{exc}}$ | 3.0 | ms | Gerstner *et al.* (2014) |
| $\tau_{\text{inh}}$ | 5.0 | ms | Gerstner *et al.* (2014) |
| $N_{\text{syn}}$ | 5 000 | - | Goetz *et al.* (2021) |
| $N_{\text{exc}}$ | 3 750 | - | Goetz *et al.* (2021) |
| $N_{\text{inh}}$ | 1 250 | - | Goetz *et al.* (2021) |
| $\mu_{w_{\text{exc}}}$ | 2.07 | - | Padamsey *et al.* (2022) |
| $\sigma_{w_{\text{exc}}}$ | 0.64 | - | Padamsey *et al.* (2022) |
| $\mu_{w_{\text{inh}}}$ | 2.07 | - | Padamsey *et al.* (2022) |
| $\sigma_{w_{\text{inh}}}$ | 0.64 | - | Padamsey *et al.* (2022) |
| $\mu_{r_{\text{exc}}}$ | 1.37 | Hz | Goetz *et al.* (2021) |
| $\sigma_{r_{\text{exc}}}$ | 0.23 | Hz | Goetz *et al.* (2021) |
| $\mu_{r_{\text{inh}}}$ | 6 | Hz | Goetz *et al.* (2021) |
| $\sigma_{r_{\text{inh}}}$ | 0 | Hz | Goetz *et al.* (2021) |

**Table 3:** Parameters used to estimate the neuronal energy budget. Morphological and physiological values approximate a layer 2/3 rat pyramidal neuron.

| Parameter | Value | Unit | Reference |
| --- | --- | --- | --- |
| $E_{HK}$ | $5.38 \cdot 10^8$ | ATP/s | Yu *et al.* (2017) |
| $E_{Na^+}$ | 50 | mV | Attwell and Laughlin (2001) |
| $E_{K^+}$ | -100 | mV | Attwell and Laughlin (2001) |
| $e$ | $1.6 \cdot 10^{-19}$ | C | - |
| $A_{overlap}$ | 1.24 | - | Carter and Bean (2009) |
| $d_{axon}$ | 0.3 | $\mu m$ | Braitenberg and Schüz (1998) |
| $l_{axon}$ | 4 | cm | Braitenberg and Schüz (1998) |
| $d_{soma}$ | 25 | $\mu m$ | Attwell and Laughlin (2001) |
| $d_{dendrite}$ | $3 \cdot d_{axon}$ | $\mu m$ | Braitenberg and Schüz (1998) |
| $l_{dendrite}$ | $l_{axon}/9$ | cm | Braitenberg and Schüz (1998) |
| $c_m$ | 0.01 | F/m<sup>2</sup> | Attwell and Laughlin (2001) |
| $V_{peak}$ | 25 | mV | Carter and Bean (2009) |
| $V_{th}$ | -50 | mV | Zerlaut and Destexhe (2017) |
| $A_{syn}$ | 1.1 | - | |
| $N_{boutons}$ | 8 000 | - | Braitenberg and Schüz (1998) |
| $E_{glu,rec}$ | $1.468 \cdot 10^4$ | ATP/vesicle | Yu *et al.* (2017) |
| $E_{Ca^{2+}ex}$ | $1.2 \cdot 10^4$ | ATP/vesicle | Attwell and Laughlin (2001) |

## Acknowledgments

We want to thank Zahid Padamsey (Weill Cornell Medicine, Qatar) for his provision and explanation of the experimental data, as well as his feedback on our results.

## Author contributions

P.S., A.D.B., P.J., J.T. designed the study.

P.S. performed the simulations and analyzed the data.

P.S., A.D.B., P.J., J.T. wrote the paper.

During the preparation of this work the authors used ChatGPT-5 for inspirational language work, improvements to human-generated texts for readability and style, and to ensure that the texts are free of errors in grammar, spelling, punctuation, and tone as well as the readability and debugging of code. After using this tool, the authors reviewed and edited the content as needed and take full responsibility for the content of the published article.

## Materials and methods

### Experimental data

The raw experimental data is accessible by the original publications (Padamsey *et al*., 2022; Zeldenrust *et al*., 2024). Extracted data and example data is available with the analysis code on github (https://github.com/trieschlab/energy-dependent-IF-model).

### Neuron model

We implemented leaky integrate-and-fire (LIF) (Brunel and Van Rossum, 2007) and adaptive exponential integrate-and-fire (AdExp) (Brette and Gerstner, 2005) point-neuron models and combined both with an updated neuronal energy budget (Sommer *et al*., 2026) based on Attwell and Laughlin (2001); Howarth *et al*. (2012); Yu *et al*. (2017) and the experimental findings of Padamsey *et al*. (2022). For all results presented here, we used the AdExp model.

The LIF membrane potential followed:

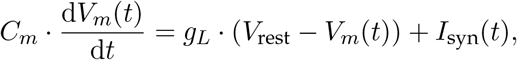

where *C_m_* is the membrane capacitance, *V_m_*(*t*) the membrane potential, *g_L_* the leak conductance, *V*_rest_ the leak reversal potential, and *I*_syn_(*t*) the synaptic input current. Whenever *V_m_*(*t*) exceeded the spike threshold *V*_th_, a spike was recorded and the membrane potential was set to *V*_reset_.

To reproduce neuronal behavior more adequately, we used the AdExp model (Brette and Gerstner, 2005) which extends the basic LIF model by adding an exponential spike-initiation term and an adaptation current, thereby capturing both rapid spike onset and activity-dependent changes in excitability. The membrane potential dynamics were given by:

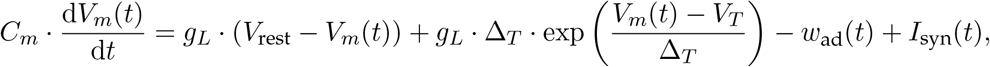

and the adaptation current followed:

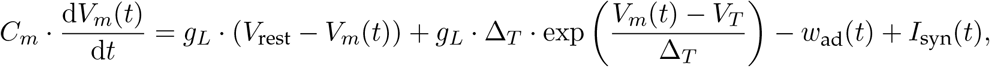

Here, Δ*_T_* is the slope factor of the exponential spike-initiation term, *V_T_* the exponential onset voltage, *w*_ad_(*t*) the adaptation current, *ε*_ad_ the adaptation time constant, and *a*_ad_ the subthreshold adaptation conductance. Whenever *V_m_*(*t*) reached the spike threshold, a spike was registered, the membrane potential was reset, and the adaptation current *w*_ad_(*t*) was increased by the spike-triggered adaptation parameter *b*_ad_.

#### Stochastic membrane noise

To capture intrinsic subthreshold fluctuations, we added a stochastic noise term to the membrane dynamics. Noise was modeled as temporally uncorrelated Gaussian input with zero mean and amplitude σ_noise_, which was estimated from experimental *ex vivo* mEPSC recordings (Padamsey *et al*., 2022; Sommer *et al*., 2026). In the Euler update, the stochastic increment was scaled by Δ*t*,

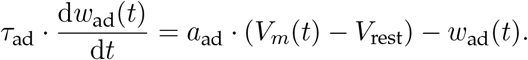

where *N* (0, 1) denotes a standard normal random variable and Δ*t* is the integration timestep. Simulations containing stochastic input were integrated using the forward Euler method, whereas noise-free simulations were performed using the deterministic Runge–Kutta solver implemented in Brian2.

### Synaptic input model

The total synaptic input current was modeled as the sum of excitatory and inhibitory conductance-based synaptic currents:

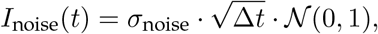

with:

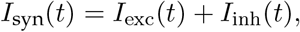

where *g*_exc_(*t*) and *g*_inh_(*t*) denote the excitatory and inhibitory synaptic conductances, and *E*_exc_ and *E*_inh_ their respective reversal potentials. Synaptic conductances decayed exponentially according to:

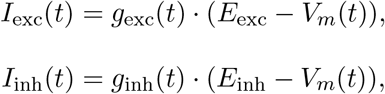

where *ε*_exc_ and *ε*_inh_ are the excitatory and inhibitory conductance decay time constants. Each presynaptic spike increased the corresponding conductance by the synaptic weight *w*exc or *w*inh.

To reproduce the experimentally observed heterogeneity of excitatory inputs, we sampled excitatory weights from a log-normal distribution fitted to the *ex vivo* mEPSC amplitudes measured in control animals, using *µ_w_*exc = 2.07 and *ϖ_w_*exc = 0.64 (Padamsey *et al*., 2022). This approach treats mEPSC amplitude as an experimental proxy for synaptic efficacy and is consistent with evidence that biological synaptic strengths are approximately log-normally distributed (Song *et al*., 2005). Inhibitory weights were sampled from the same underlying distribution and subsequently rescaled so that their summed weight equaled 25 % of the summed excitatory weight, yielding an overall excitation–inhibition balance (Deneve and Machens, 2016). A shared scaling parameter, *w*_scale_, was then applied to both excitatory and inhibitory weights. This preserved their relative balance while allowing the total strength of synaptic input to be varied in accordance with the proportional changes reported experimentally (Padamsey *et al*., 2022). At *w*_scale_ = 100, the resulting mean weights were *w*_exc_ = 0.52 nS and *w*_inh_ = 0.40 nS, respectively.

Presynaptic input activity was generated by independent homogeneous Poisson spike trains. Excitatory presynaptic firing rates were drawn from a normal distribution with mean *µ_r_*exc = 1.37 Hz and standard deviation *ϖ_r_*exc = 0.23 Hz (Goetz *et al*., 2021). All inhibitory inputs fired at 6 Hz. Each postsynaptic model neuron received 5 000 synaptic inputs in total, comprising 3 750 excitatory and 1 250 inhibitory synapses.

### Updated energy consumption budget

Neuronal energy consumption was estimated using the updated neuronal energy budget derived in Sommer *et al*. (2026), which extends the framework of Attwell and Laughlin (2001). Here, we briefly summarize the components required to interpret the simulations.

#### Housekeeping energy demand

Neurons consume energy even in the absence of spiking or synaptic signaling. This activity-independent demand includes macromolecule synthesis and turnover, cytoskeletal remodeling, axoplasmic transport, mitochondrial proton leak, and other nonsignaling processes (Engl and Attwell, 2015; Balasubramanian, 2021). We grouped these baseline costs into a housekeeping term, *E*_HK_, defined as the number of ATP molecules hydrolyzed per second to maintain basic cellular function. Following Yu *et al*. (2017), we treated *E*_HK_ as constant across activity regimes.

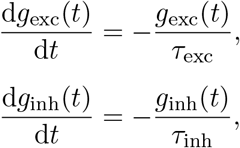

#### Resting potential energy demand

Maintaining the resting membrane potential requires continuous ATP-dependent ion pumping to compensate passive ionic leak. We estimated this cost as *E*_RP_, the number of ATP molecules hydrolyzed per second to maintain a neuron at a resting potential *V*_RP_. Following Attwell and Laughlin (2001), this estimate assumes that resting ionic fluxes are balanced by the sodium–potassium pump, which extrudes three Na^+^ ions and imports two K^+^ ions per ATP molecule. To account for HCN-mediated resting currents, we extended the original sodium–potassium leak estimate by an additional *I_h_*term:

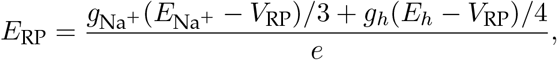

where *g*Na+ and *g_h_* denote the sodium and HCN conductances, *E*Na+ and *E*_h_ their reversal potentials, and *e* the elementary charge.

#### Action potential energy demand

Action potentials consume energy because the sodium influx that depolarizes the membrane must subsequently be extruded by the sodium–potassium pump. We estimated the spike-related energy demand, *E*_AP_, from the sodium charge entering during one action potential and scaled this cost by the mean postsynaptic firing rate, *r*_post_:

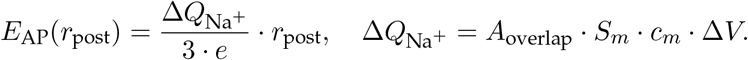

Here, *e* is the elementary charge, *S_m_* is the total membrane surface area, *c_m_* is the specific membrane capacitance, and Δ*V* is the voltage difference between spike threshold and spike peak. The factor *A*_overlap_ accounts for temporal overlap between sodium influx and potassium efflux, which increases sodium entry beyond the theoretical capacitive minimum (Alle *et al*., 2009; Carter and Bean, 2009).

#### Synaptic transmission energy demand

Excitatory synaptic transmission consumes postsynaptic energy because ionotropic receptor activation drives inward ionic currents whose gradients must be restored by ATP-dependent pumps. We estimated this cost from the total excitatory synaptic current entering the postsynaptic neuron:

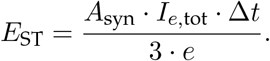

Here, organized sensory circuits (Van Essen and Maunsell, 1983).

#### Presynaptic neurotransmitter recycling and calcium extrusion

Presynaptic signaling consumes energy through neurotransmitter recycling, vesicle refilling, and calcium extrusion after vesicle release (Li and Sheng, 2022). We modeled these presynaptic costs as activity-dependent terms that scale with the output firing rate, the release probability, and the number of downstream boutons:

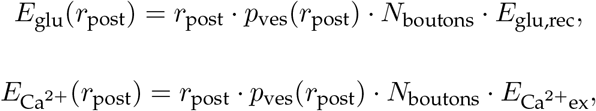

with

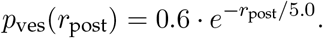

Here, and Markram, 1997; Tsodyks *et al*., 1998; Bird and Richardson, 2018). Following previous estimates, one glutamate vesicle was assumed to contain approximately 4000 glutamate molecules, with 3.67 ATP molecules required per recycled glutamate molecule (Riveros *et al*., 1986; Yu *et al*., 2017), whereas calcium extrusion was estimated from 12 000 Ca^2+^ ions per released vesicle and one ATP molecule per extruded Ca^2+^ ion (Attwell and Laughlin, 2001). Minor energetic contributions from inhibitory chloride currents and other ionic species were not modeled as separate terms but were included in the synaptic correction factor *A*_syn_.

### Internal state variables

To coordinate energy demand with stable neuronal output, parameter adaptation was driven by two state estimates: a spike-derived activity signal and an energy balance. The former provides feedback to maintain firing-rate homeostasis, whereas the latter constrains consumption to the available metabolic budget.

#### Neuronal activity variable

Recent spiking activity was estimated using a spike-triggered calcium proxy, motivated by the capacity of intracellular Ca^2+^-dependent pathways to integrate spiking over time (LeMasson *et al*., 1993; Helmchen *et al*., 1996). Each spike incremented *r*Ca2+ (*t*), which otherwise decayed exponentially:

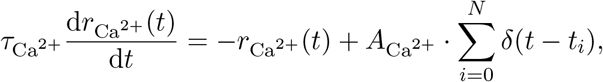

where *T*_Ca_^2+^ is the decay time constant, *A*_Ca_^2+^ is the spike-triggered calcium increment, and *t_i_* denotes the spike times. This calcium-like trace provided a phenomenological readout of the activity-dependent intracellular signaling pathways involved in homeostatic regulation (LeMasson *et al*., 1993; Helmchen *et al*., 1996; Marder and Prinz, 2002; O’Leary *et al*., 2013).

#### Energy balance variable

In parallel, the energetic state of the neuron was represented by a metabolic feedback variable. Biologically, cellular energy state can be sensed by AMP-activated protein kinase (AMPK), which is regulated by the AMP/ATP ratio and acts as a central mediator of energy homeostasis (Potter *et al*., 2010). Motivated by AMP-activated protein kinase (AMPK), which responds to cellular AMP/ATP balance, thus acting as a central mediator of energy homeostasis (Potter *et al*., 2010), we defined this signal as the ratio of available ATP production to instantaneous consumption:

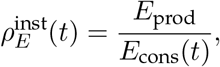

where *E*_cons_(*t*) is the instantaneous ATP consumption estimated from the energy budget, and *E*_prod_ is the available ATP production. The balanced operating point was defined as *ω_E,_*_target_ = 1, at which ATP production equals consumption.

Consistent with supply-limited accounts of neuronal metabolism, available ATP production was bounded by a target energy budget (Herculano-Houzel, 2011; Herculano-Houzel and Rothman, 2022):

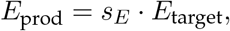

where *E*_target_ is the target energy budget and *s_E_*is an energy-supply fraction representing the metabolic condition of the animal. We used *s_E_* = 1 for control and *s_E_* = 0.85 for food restriction, representing an estimated 15 % reduction in available energy (Padamsey *et al*., 2022, 2024). Thus, ATP production was treated as fixed over the adaptation time scale, whereas ATP consumption was dynamically regulated to approach the imposed supply.

#### Low-pass filtered feedback signals

Because the experimental phenotype develops over days to weeks, the control rules should respond to sustained rather than momentary deviations in activity and energy balance (Padamsey *et al*., 2022). We therefore low-pass filtered both state estimates before they entered the parameter-adaptation rules:

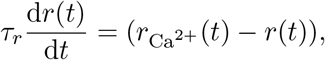

and:

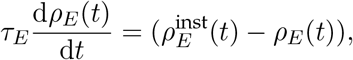

where *r*(*t*) is the slow activity estimate and *ω_E_*(*t*) is the slow energy-balance estimate.

## Supporting information

### Frequently used abbreviations

- AdExp — Adaptive exponential integrate-and-fire neuron model
- ATP — Adenosine triphosphate
- CTR — Control condition; normally fed mice
- CV — Coefficient of variation
- FR — Food-restricted condition; mice subjected to 2 weeks of dietary restriction
- ISI — Inter-spike interval
- LIF — Leaky integrate-and-fire neuron model

### 1 Proportional control alone results in oscillatory parameter dynamics

To determine whether proportional feedback was sufficient for firing-rate homeostasis, we repeated the simulation without the derivative component of the intrinsic controller. In this configuration, the slow firing-rate estimate repeatedly crossed its target because *V*_rest_ and *R_m_* continued to adapt while the activity signal lagged behind the changing output (Fig. S1). The resulting oscillations demonstrate that proportional control alone responds too slowly to prevent repeated overshooting. We therefore retained the derivative term in the main model to oppose rapid changes in activity and stabilize convergence.

**Figure S1:**
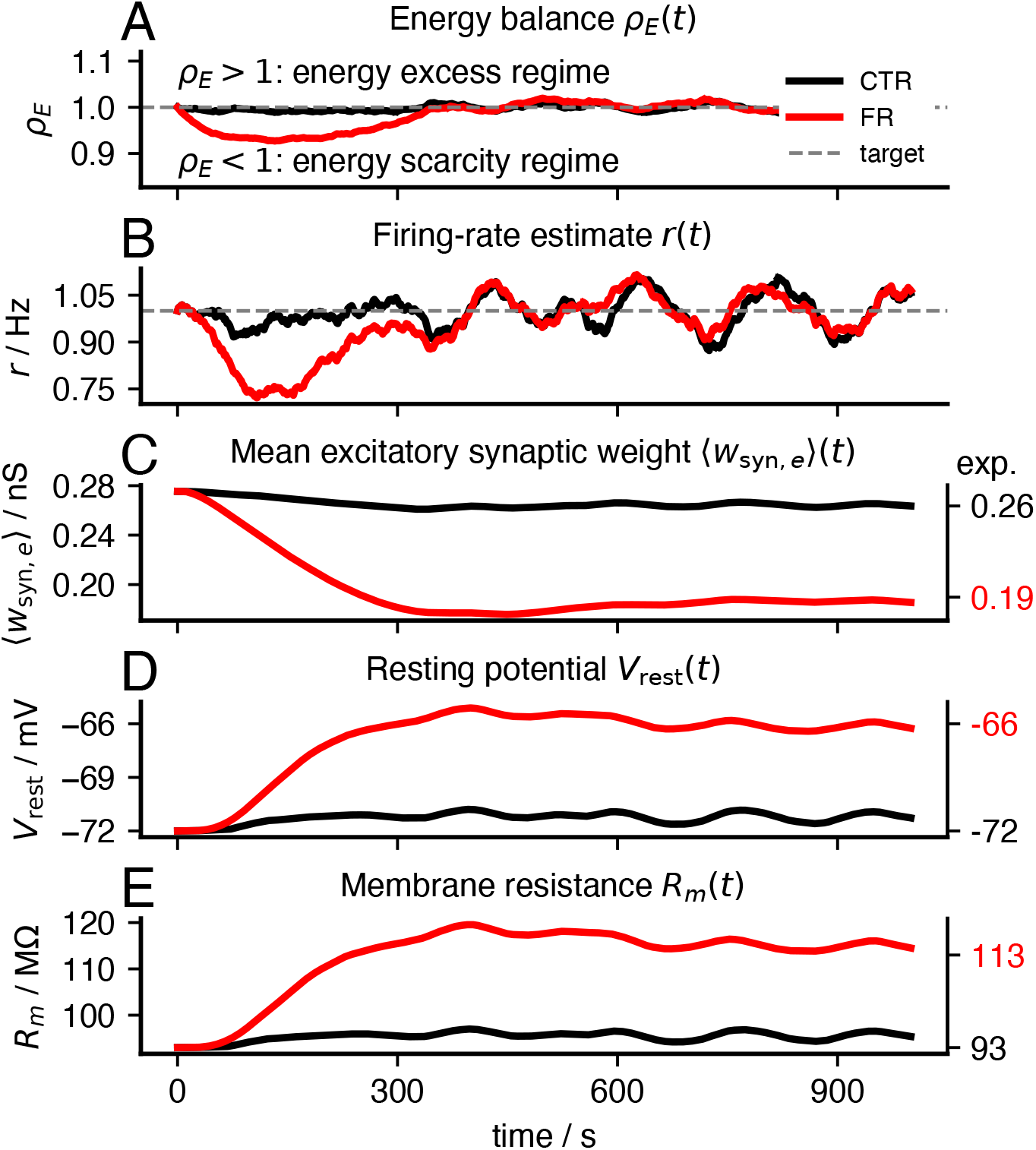
Proportional control alone produces oscillatory firing-rate and intrinsic-parameter dynamics. Simulation of the energy-dependent integrate-and-fire model with only proportional control in the intrinsic plasticity rule. **A**, Energy-balance signal *ω_E_*(*t*) under normal energy supply (control, CTR, black; *s_E_* = 1) and reduced energy supply (food restriction, FR, red; *s_E_*= 0.85). **B**, Firing-rate estimate *r*(*t*). Without derivative damping, the firing-rate estimate oscillates around the target firing rate. **C**, Mean excitatory synaptic weight *w*_syn,e_ (*t*). Energy-dependent synaptic scaling still reduces synaptic strength under reduced energy supply. **D–E**, Resting potential and membrane resistance. Proportional control alone causes *V*_rest_ and *R_m_* to overshoot the target regime, producing oscillatory intrinsic-parameter dynamics. This instability motivates the derivative control term used in the main model.

### 2 Blocking resting-potential adaptation shifts compensation onto membrane resistance

To determine whether the two intrinsic variables can compensate for one another, we repeated the *synapse-first* simulation while holding *V*_rest_ fixed. Energy-dependent synaptic downscaling still reduces synaptic drive and transiently suppresses firing, but activity could no longer recover through resting-potential depolarization. *R_m_* therefore increases more strongly, indicating that homeostatic compensation is redirected to the remaining intrinsic degree of freedom (Fig. S2).

**Figure S2:**
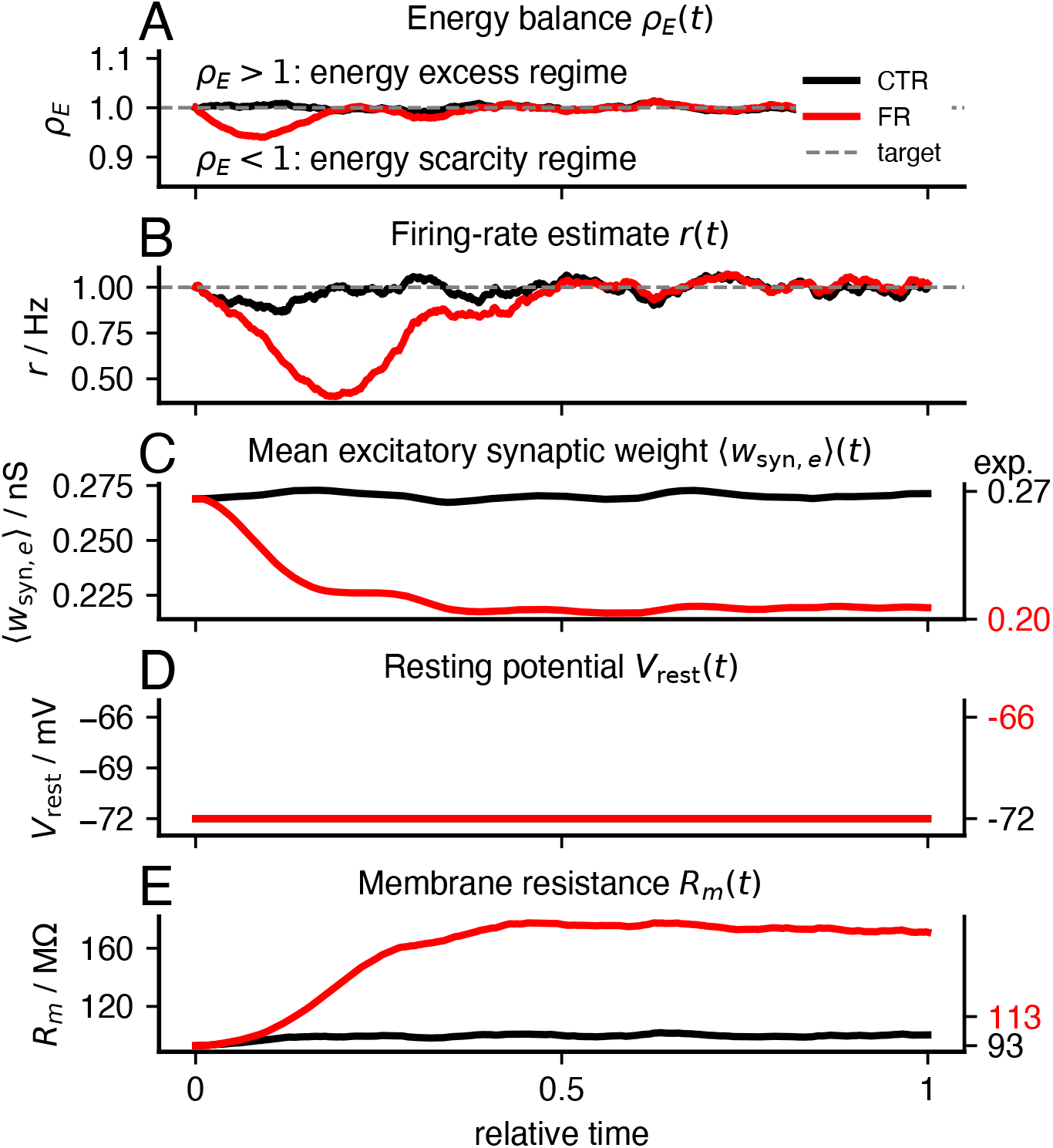
Constraining resting-potential adaptation shifts compensation onto membrane resistance. Simulation of the *synapse-first* control architecture under normal energy supply (control, CTR, black; *s_E_* = 1) and reduced energy supply (food restriction, FR, red; *s_E_* = 0.85), with resting-potential adaptation constrained. **A–C**, Reduced energy supply in the FR condition lowers the energy-balance signal *ω_E_*(*t*), induces synaptic downscaling, and transiently reduces the firing-rate estimate *r*(*t*). **D**, Resting potential *V*_rest_ remains fixed throughout the simulation. **E**, In the absence of resting-potential plasticity, *R_m_* increases more strongly to offset the reduction in synaptic drive.

#### Energy-dependent synaptic upscaling following neuronal silencing

To investigate whether energy-dependent feedback could also explain synaptic upscaling following activity deprivation, we simulated neuronal silencing analogous to tetrodotoxin (TTX) treatment. Reduced neuronal activity decreases ATP consumption, increasing the energy-balance signal *ω_E_* and thereby promoting synaptic upscaling. At the same time, activity-dependent feedback increases membrane resistance, while resting potential is held constant, consistent with experimental observations (Wen *et al*., 2025). These results qualitatively reproduce adaptations reported following prolonged TTX treatment (Fig. S3) and suggest that energy balance may contribute to bidirectional synaptic scaling.

**Figure S3:**
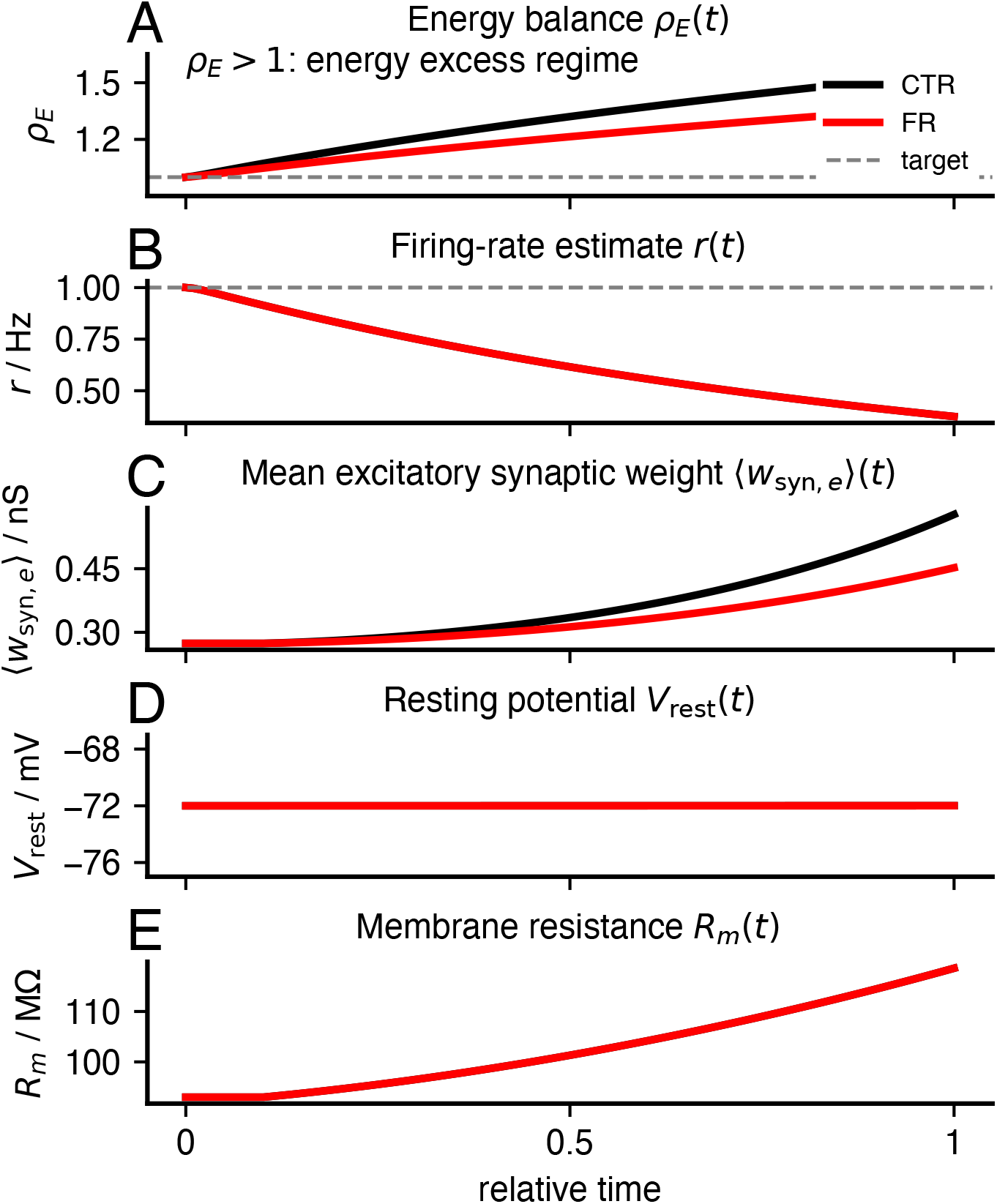
Neuronal silencing induces energy-dependent synaptic upscaling and increased membrane resistance. Simulation of the *synapse-first* control architecture under normal activity and simulated neuronal silencing, mimicking tetrodotoxin (TTX) treatment, with resting-potential adaptation constrained. **A–C**, Reduced neuronal activity decreases ATP consumption, increases the energy-balance signal *ω_E_*(*t*), and induces synaptic upscaling. **D**, Resting potential *V*_rest_ remains fixed throughout the simulation. **E**, Membrane resistance *R_m_* increases in response to reduced firing activity. Together, these adaptations qualitatively reproduce experimental observations following prolonged neuronal silencing and suggest a potential role for energy-dependent feedback in bidirectional synaptic scaling.

### 3 Empirical constraints on relative intrinsic adaptation time scales

Reproducing the food-restriction phenotype required specifying how intrinsic compensation was divided between *V*_rest_ and *R_m_*. Both variables increase excitability when firing falls below target, but their effects differ in magnitude. We therefore constrained their relative adaptation rates using the steady-state shifts measured between neurons from control (CTR) and food-restricted (FR) animals (Padamsey *et al*., 2022):

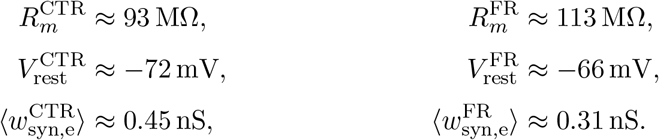

Food restriction was therefore associated with a decrease in excitatory synaptic strength:

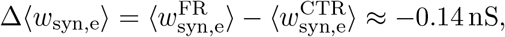

together with an increase in membrane resistance and a depolarization of the resting potential:

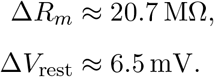

To quantify how the two intrinsic shifts accompanied synaptic downscaling, we related the change in synaptic strength to the corresponding change in each intrinsic parameter:

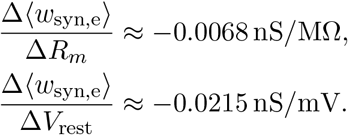

The slope associated with *V*_rest_ was approximately threefold larger than that associated with *R_m_* in absolute units. Scaling both effects by their typical physiological values, *R_m_↗* 93 MΛ and *V*_rest_ *↗ →*72 mV, yielded a comparable ratio, with the resting-potential contribution remaining approximately 2.5 times larger.

We used this ratio to constrain the relative, but not necessarily the absolute, adaptation time scales of *V*_rest_ and *R_m_*. The overall speed of plasticity therefore remained free, whereas the empirical constraint determined how compensation was partitioned between the two intrinsic variables. Under this calibration, the model approached the observed CTR–FR steady-state shift.

As an alternative constraint, the relative adaptation ratios can be derived from the geometry of the high-coding-efficiency surface identified in the parameter-space analysis performed in *Sommer et al.* (2026) (2026) (Fig. S4). A plane fitted to the region of high *OSI* per energy provided an independent local constraint:

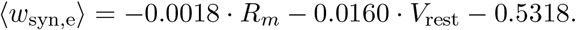

Its partial slopes were:

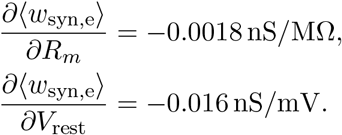

Relative to the experimental CTR–FR shift, this geometric estimate assigned greater leverage to *V*_rest_: its slope was approximately ninefold larger in absolute units and sixfold larger after normalization by representative physiological ranges.

The two calibrations address different questions. The empirical ratio describes the intrinsic adjustments accompanying the measured food-restriction phenotype and was therefore used for the primary simulations. By contrast, the plane-based ratio characterizes the local orientation of the high-efficiency manifold and may be more informative when extrapolating beyond the experimentally sampled metabolic regime.

**Figure S4:**
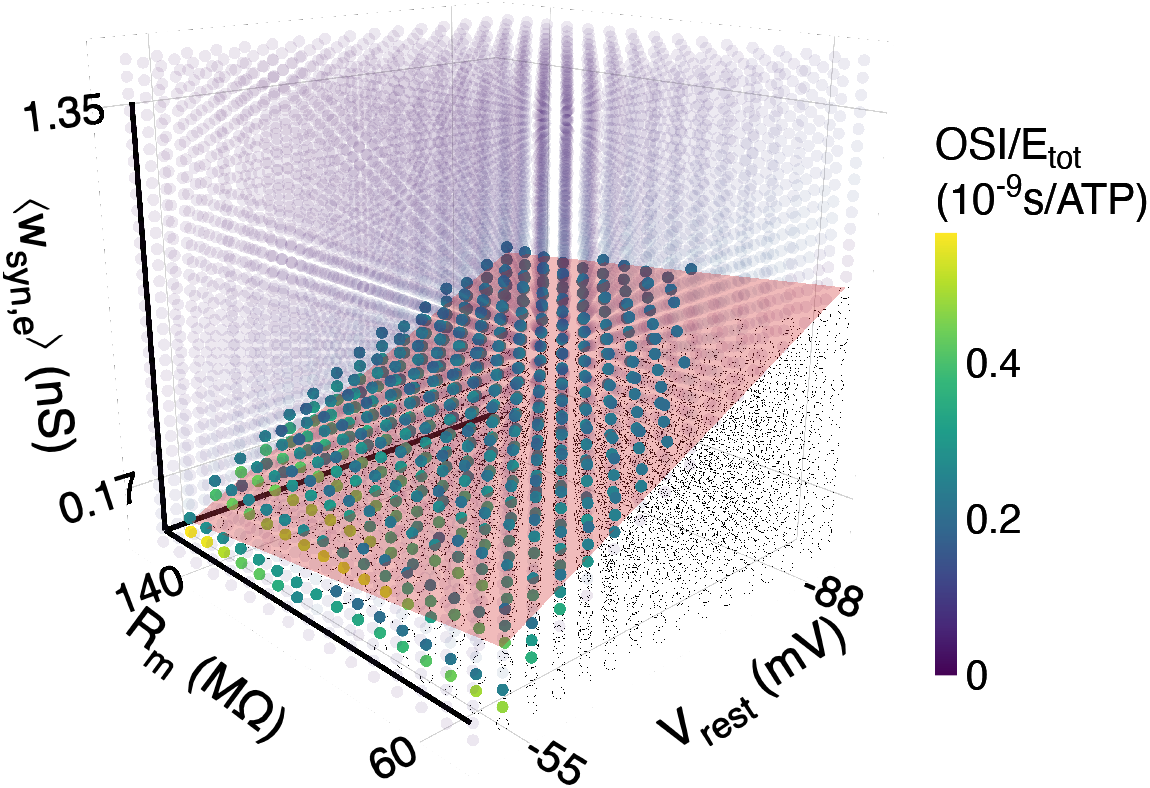
Fitted high-efficiency plane in neuronal parameter space. Plane fitted to parameter combinations with high orientation selectivity per unit energy (*OSI*/*E*_tot_) in the three-dimensional parameter space spanned by resting potential *V*_rest_, membrane resistance *R_m_*, and mean excitatory synaptic weight *w*_syn,e_. Colored points indicate simulated parameter combinations, with color denoting *OSI*/*E*_tot_. The fitted plane approximates the low-dimensional region in which neurons achieve high coding efficiency. The slopes of the plane was used as an alternative constraint for deriving relative adaptation time scales of *V*_rest_ and *R_m_* in the energy-dependent integrate-and-fire model.

